# Transformer-based spatial-temporal detection of apoptotic cell death in live-cell imaging

**DOI:** 10.1101/2022.11.23.517318

**Authors:** Alain Pulfer, Diego Ulisse Pizzagalli, Paolo Armando Gagliardi, Lucien Hinderling, Paul Lopez, Romaniya Zayats, Pau Carrillo-Barberà, Paola Antonello, Miguel Palomino-Segura, Benjamin Grädel, Mariaclaudia Nicolai, Alessandro Giusti, Marcus Thelen, Luca Maria Gambardella, Thomas T. Murooka, Olivier Pertz, Rolf Krause, Santiago Fernandez Gonzalez

**Author notes:** Contributed equally (Alain Pulfer, Diego Ulisse Pizzagalli, Paolo Armando Gagliardi).

## Abstract

Intravital microscopy has revolutionized live cell imaging by allowing the study of spatial-temporal cell dynamics in living animals. However, the complexity of the data generated by this technology has limited the development of effective computational tools to identify and quantify cell processes. Amongst them, apoptosis is a crucial form of regulated cell death involved in tissue homeostasis and host defense. Live-cell imaging enabled the study of apoptosis at the cellular level, enhancing our understanding of its spatial-temporal regulation. However, at present, no computational method can deliver robust detection of apoptosis in microscopy time-lapses. To overcome this limitation, we developed ADeS, a deep learning-based apoptosis detection system that employs the principle of activity recognition. We trained ADeS on extensive datasets containing more than 10,000 apoptotic instances collected both in vitro and in vivo, achieving a classification accuracy above 98% and outperforming state-of-the-art solutions. ADeS is the first method capable of detecting the location and duration of multiple apoptotic events in full microscopy time-lapses, surpassing human performance in the same task. We demonstrated the effectiveness and robustness of ADeS across various imaging modalities, cell types, and staining techniques. Finally, we employed ADeS to quantify cell survival in vitro and tissue damage in vivo, demonstrating its potential application in toxicity assays, treatment evaluation, and inflammatory dynamics. Our findings suggest that ADeS is a valuable tool for the accurate detection and quantification of apoptosis in live-cell imaging and, in particular, intravital microscopy data, providing insights into the complex spatial-temporal regulation of this process.

## 1. Introduction

In the last two decades, Intravital microscopy (IVM) has revolutionized live-cell imaging by enabling microscopy acquisitions *in situ* across different organs, making it one of the most accurate model to describe cellular activities within a living host(Sumen et al. 2004). In particular, multiphoton intravital microscopy (MP-IVM) generates in-depth 3D data that encompass multiple channels for up to several hours of acquisition (x,y,z + t)(Helmchen and Denk 2005; Rocheleau and Piston 2003; Secklehner, Celso, and Carlin 2017), thus providing unprecedented insights into cellular dynamics and interactions(Pizzagalli et al. 2019). The resulting MP-IVM data stream is a complex and invaluable source of information, contributing to enhance our understanding of several fundamental processes(Beltman et al. 2009; Sumen et al. 2004).

Apoptosis is a form of regulated cell death(D’Arcy 2019; D. Tang et al. 2019) which plays a crucial role in several biological functions, including tissue homeostasis, host protection, and immune response(Opferman 2008). This process relies on the proteolytic activation of caspase-3-like effectors(Shalini et al. 2015), which yields successive morphological changes that include cell shrinkage, chromatin condensation, DNA fragmentation, membrane blebbing(Elmore 2007; Galluzzi et al. 2018; Saraste and Pulkki 2000), and finally, apoptotic bodies formation(Coleman et al. 2001). Due to its crucial role, dysregulations of apoptosis can lead to severe pathological conditions, including chronic inflammatory diseases and cancer(Fesik 2005; Hotchkiss and Nicholson 2006). Consequently, precise tools to identify and quantify apoptosis in different tissues are pivotal to gain insights on this mechanism and its implications at the organism level.

Traditional techniques to quantify apoptosis rely on cellular staining on fixed cultures and tissues(Atale et al. 2014; Kyrylkova et al. 2012; Loo 2011; Sun et al. 2008; Vermes et al. 1995) or flow cytometry(Darzynkiewicz, Galkowski, and Zhao 2008; Vermes et al. 1995). However, these methods do not allow the temporal characterization of the apoptotic process. Moreover, they potentially introduce artifacts caused by sample fixation(Schnell et al. 2012). Live-cell imaging can overcome these limitations by unraveling the dynamic aspects of apoptosis with the aid of fluorescent reporters, such as Annexin staining(Atale et al. 2014) or the activation of Caspases(Takemoto et al. 2003). However, the use of fluorescent probes *in vivo* could potentially interfere with physiological functions or lead to cell toxicity(Jensen 2012). For these reasons, probe-free detection of apoptosis represent a critical advancement in the field of cell death.

Computational methods could address this need by automatically detecting individual apoptotic cells with high spatial and temporal accuracy. In this matter, deep learning (DL) and activity recognition (AR) could provide a playground for the classification and detection of apoptosis based on morphological features(Poppe 2010). Accordingly, recent studies showed promising results regarding the classification of static frames(Kranich et al. 2020; Verduijn et al. 2021) or time-lapses(Mobiny et al. 2020) portraying single apopotic cells. However, none of the available methods can be applied for the detection of apoptosis in microscopy movies depicting multiple cells. Therefore we developed ADeS, a novel apoptosis detection system which employs a transformer DL architecture and computes the location and duration of multiple apoptotic events in live-cell imaging. Here we show that our architecture outperforms state-of-the-art DL techniques and efficiently detects apoptotic events in a broad range of imaging modalities, cellular staining, and cell types.

## 2. Results

### 2.1. An *in vitro* and *in vivo* live-cell imaging data

Curated and high-quality datasets containing numerous instances of training samples are critical for developing data-hungry methods such as supervised DL algorithms(Adadi 2021). To this end, we generated two distinct datasets encompassing epithelial cells (*in vitro*) and leukocytes (*in vivo*) undergoing apoptotic cell death. In addition, the two datasets include different imaging modalities (confocal and intravital 2-photon), biological models, and training-set dimensionalities. A meaningful difference between the datasets pertains to the staining methods and the morphological hallmarks, which define the apoptotic process in both models. In the *in vitro* model, the expression of nuclear markers allowed us to observe apoptotic features such as chromatin condensation and nuclear shrinkage(Saraste and Pulkki 2000), whereas in the *in vivo* model, cytoplasmic and membrane staining highlighted morphological changes such as membrane blebbing and the formation of apoptotic bodies(Saraste and Pulkki 2000). Accordingly, we have manually annotated these datasets based on the presence of the specific hallmarks, ensuring that each dataset includes two class labels depicting either apoptotic or non-apoptotic cells. These two datasets constitute the first step toward creating, testing, and validating our proposed apoptosis detection routine.

To generate the *in vitro* dataset we used epithelial cells because, among the human tissues, they have the highest cellular turnover driven by apoptosis(Van Der Flier and Clevers 2009). Nevertheless, from the bioimaging perspective, the epithelium is a densely packed tissue with almost no extracellular matrix, making it extremely challenging to analyze. As such, in epithelial research, there is a pressing need for computational tools to identify apoptotic events automatically. To this end, we imaged and annotated the human mammary epithelial cells expressing a nuclear fluorescent marker (Fig.1A), obtaining 13120 apoptotic nuclei and 301630 non apoptotic nuclei image sequences (Fig. 1B-C, Supplementary 1A). Nuclear shrinkage and chromatin condensation, two of the most prototypical hallmarks of apoptosis (Fig. 1C), formed our criteria for manual annotation. We confirmed that non-apoptotic nuclei had constant area and chromatin density from the generated time-lapses. In contrast, apoptotic nuclei underwent a decrease in area and an increase in chromatin condensation (Fig. 1D). The resulting dataset captured the heterogeneity of apoptotic cells in epithelial tissue, including early nuclear fragmentation, a rapid shift along the x and y axes, and extrusion through the z dimension (Supplementary Fig. 1B–C). Moreover, our dataset incorporates the typical difficulties of automatically annotating apoptotic events from live microscopy of a densely packed tissue (Supplementary Fig. 1D) with the accumulation of apoptotic bodies (Supplementary Fig. 1E) and across multiple microscope hardware settings (Supplementary Fig. 1F).

**Figure 1:**
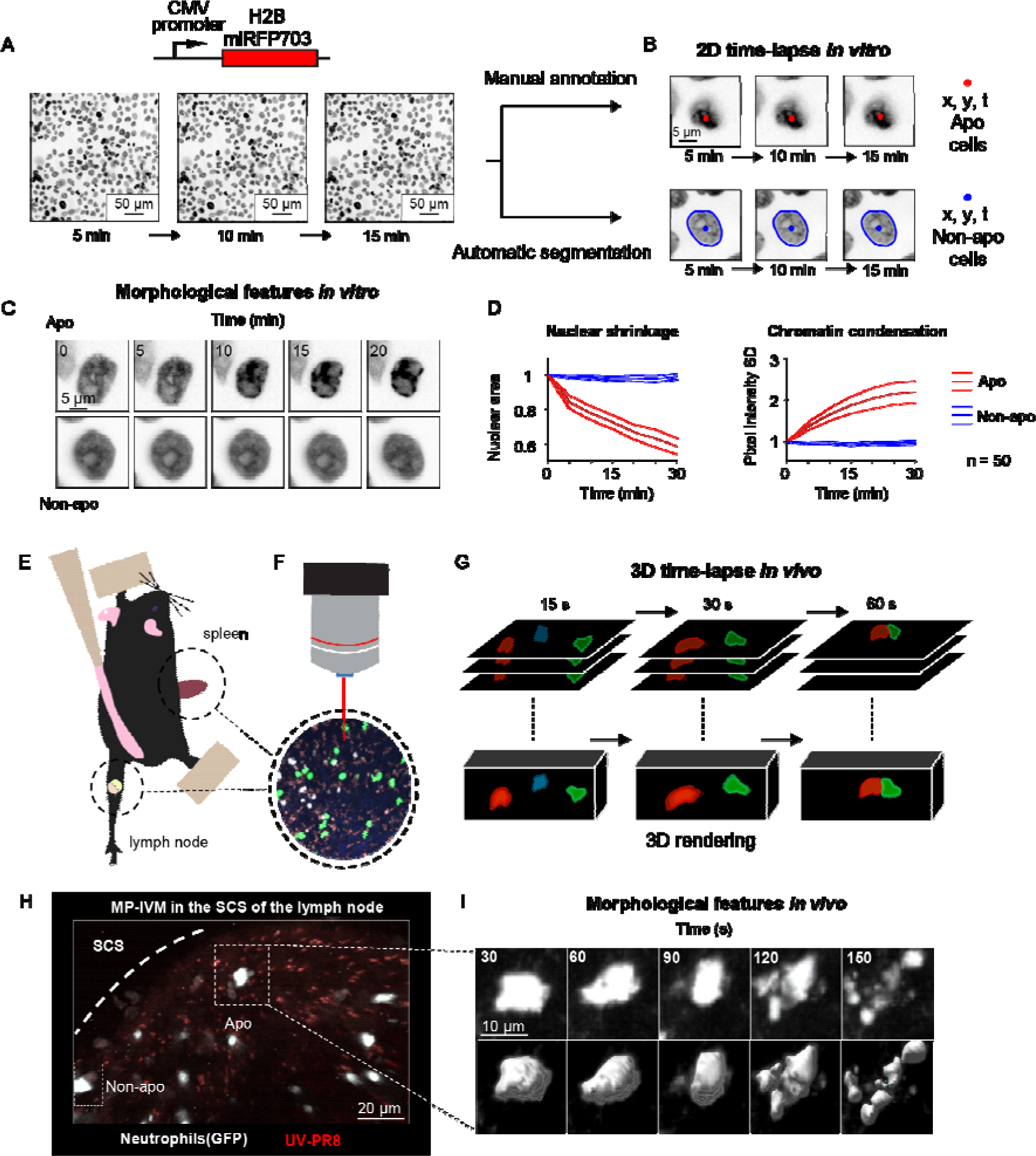
Generation of in vitro and in vivo live-cell imaging data. **A.** Micrographs depicting mammary epithelial MCF10A cells transduced with H2B-miRFP703 marker and grown to form a confluent monolayer. The monolayer was acquired with a fluorescence microscope for several hours with 1-, 2- or 5-min time resolution. **B.** The centroid (x, y) and the time (t) of apoptotic events were annotated manually based on morphological features associated with apoptosis. Non-apoptotic cells were identified by automatic segmentation of nuclei. **C.** Image time-lapses showing a prototypical apoptotic event (**upper panels**), with nuclear shrinkage and chromatin condensation, and a non-apoptotic event (**bottom panels**). **D.** Charts showing the quantification of nuclear size (**left**) and the standard deviation (SD) of the nuclear pixel intensity (**right**) of apoptotic and non-apoptotic cells (n = 50). Central darker lines represent the mean, gray shades bordered by light colored lines represent the standard deviation. Nuclear area over time expressed as the ratio between areas at Tn and T0. **E.** Simplified drawing showing the surgical set-up for lymph node and spleen. **F-G**. Organs are subsequently imaged with intravital 2-photon microscopy (IV-2PM, **F**), generating 3D time-lapses (**G**). **H.** Representative IV-2PM micrograph and **I.** selected crops showing GFP-expressing neutrophils (white) undergoing apoptosis. The apoptosis sequence is depicted by raw intensity signal (**upper panels**) and 3D surface reconstruction (**bottom panels**).

To generate an *in vivo* dataset, we focused on polymorphonucleated leukocytes (neutrophils and eosinophils) that expressed a fluorescent marker. In these early immune responders, apoptosis is a crucial process that orchestrates their disposal, consequently determining the duration of the inflammation(Fox et al. 2010). To acquire instances of apoptotic leukocytes, we performed MP-IVM in anesthetized mice by surgically exposing either the spleen or the popliteal lymph node (Fig. 1E-F). The resulting time-lapses (Fig. 1G) provided 3D imaging data encompassing consecutive multi-focal planes (3D) and multiple imaging channels. Then, from the generated MP-IVM movies, we generated cropped sequences of fixed size that tracked apoptotic cells for the duration of their morphological changes (59×59 pixels + time; Fig. 1H-I). This procedure was applied to 30 MP-IVM movies, generating 120 apoptotic sequences (supplementary Fig. 1G). Furthermore, we annotated random instances of non-apoptotic events, generating 535 cropped samples. To characterize the heterogeneity of the movies, we manually quantified the cell number per field of view (87 ± 76), the shortest distance between cells (21.2 μM ± 15.4), and the signal-to-noise ratio (8.9 ± 3.6; supplementary Fig. 1 H–J). We assumed that the morphological changes associated with apoptosis occur within defined time windows for detection purposes. Hence, we estimated the median duration of the morphological changes corresponding to 8 frames (supplementary Fig. 1K–L, respectively). In addition, to classify apoptotic cells within defined spatial regions, we considered them to be non-motile. This assumption was confirmed when we found that apoptotic cells, despite having a longer track-length due to passive transport, exhibited a speed that was not significantly different from those of arrested cells (supplementary Fig. 1M).

### 2.2. ADeS, a pipeline for apoptosis detection

Detecting apoptosis in live-cell imaging is a two-step process involving the correct detection of apoptotic cells in the movies (x,y) and the correct estimation of the apoptotic duration (*t*). To fulfill these requirements, we designed ADeS as a set of independent modules assigned to distinct computational tasks (Fig. 2). As an input, ADeS receives a 2D representation of the microscopy acquisitions (Fig. 2A) obtained from the normalization of 2D raw data or the maximum projection of 3D data(Shi et al. 2015). This processing step ensures the standardization of the input, which might differ in bit depth or acquisition volume. After that, we employ a selective search algorithm(Girshick 2015; Uijlings et al. 2013) to compute regions of interest (ROIs) that might contain apoptotic cells (Fig. 2B). For each ROI at time (*t*), ADeS extracts a temporal sequence of *n* frames ranging *from t* - *n*/2 to *t* + *n*/2 (Fig. 2C). The resulting ROI sequence is standardized in length and passed to a DL classifier (Fig. 3), which determines whether it is apoptotic or non-apoptotic. Finally, each apoptotic sequence is depicted as a set of bounding boxes and associated probabilities (Fig. 2D) generated from the predicted trajectories (x, y, t, ID; Fig. 2E). From this readout, ADeS can generate a heatmap representing the likelihood of apoptotic events throughout a movie (Fig. 2F, left), together with a cumulative sum of the predicted cell deaths (Fig. 2F, right).

**Figure 2.**
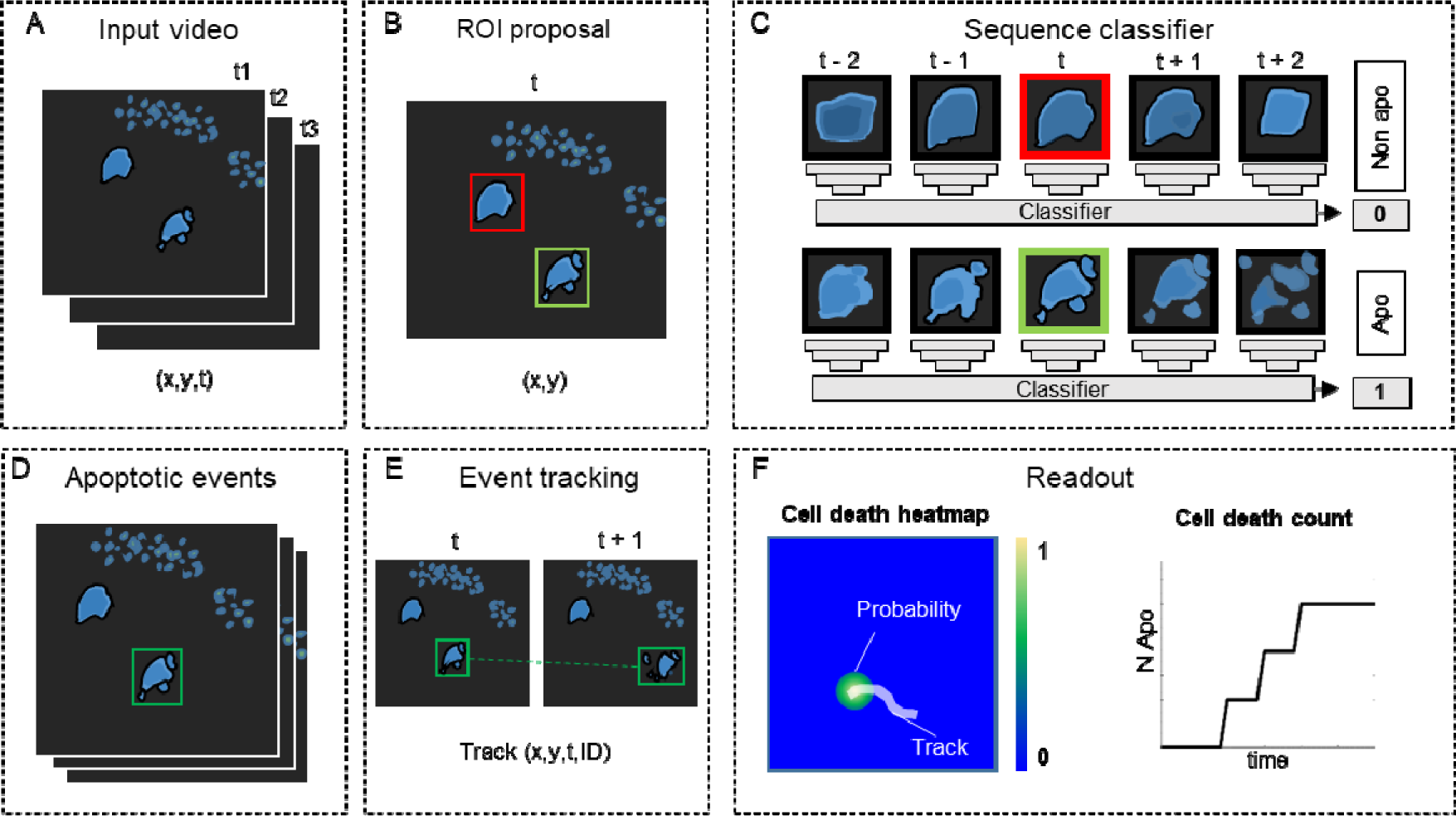
ADeS, a pipeline for apoptosis detection. **A.** ADeS input consists of single channel 2D microscopy videos (x,y,t) **B.** Each video frame is pre-processed to compute the candidate Regions of Interest (ROI) with a selective search algorithm. **C.** Given the coordinates of the ROI at time t, ADeS extracts a series of snapshots ranging from t-n to t+n. A deep learning network classifies the sequence either as non-apoptotic (0) or apoptotic (1). **D.** The predicted apoptotic events are labelled at each frame by a set of bounding boxes which, **E.** are successively linked in time with a tracking algorithm based on euclidean distance. **F.** The readout of ADeS consist of bounding boxes and associated probabilities, which can generate a probability map of apoptotic events over the course of the video (**left**) as well as providing the number of apoptotic events over time (**right**).

**Figure 3.**
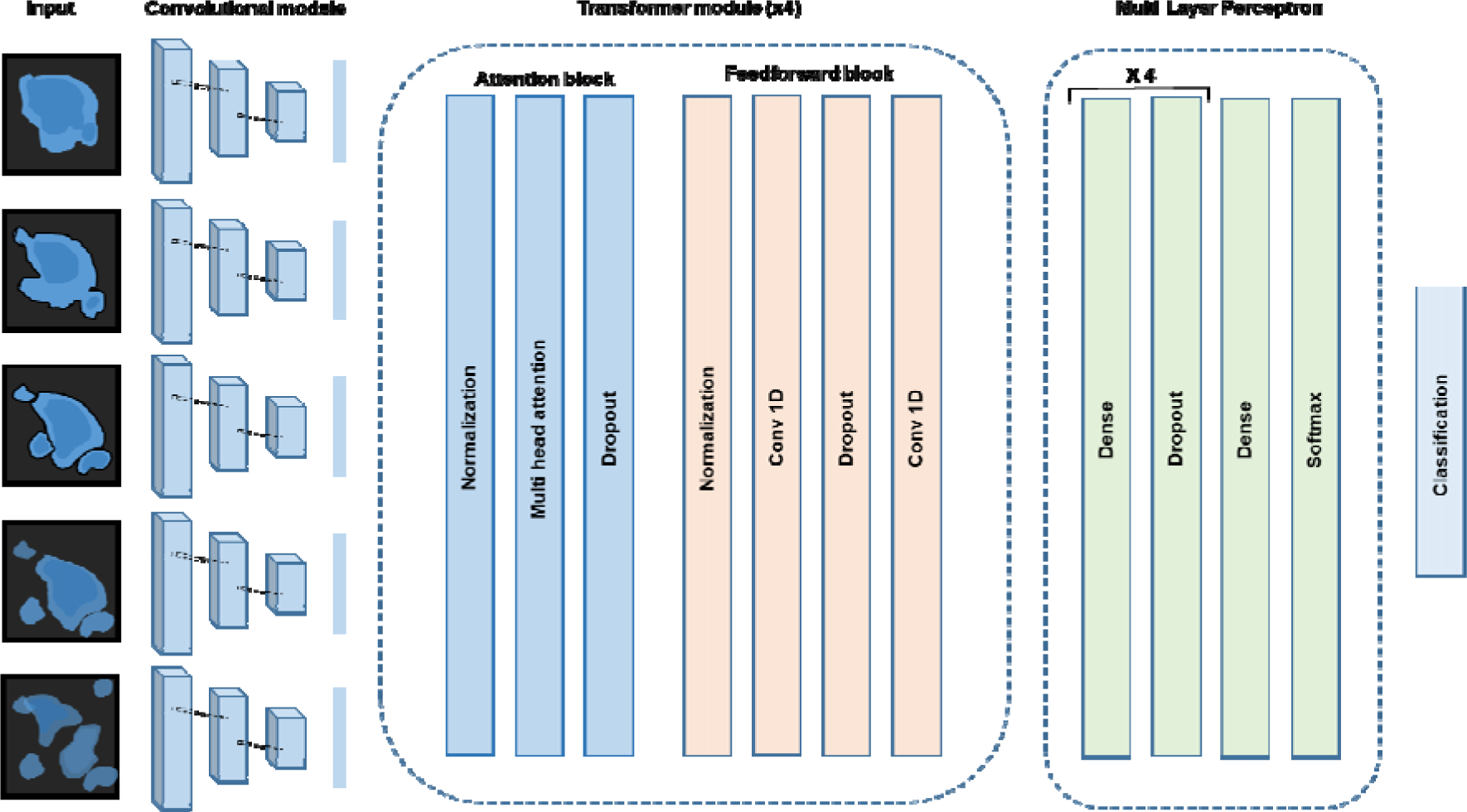
Conv-Transformer architecture at the core of ADeS. Abstracted representation of the proposed Conv-Transformer classifier. The input sequence of frames is processed with warped convolutional layers, which extract the features of the images. The extracted features are passed into the 4 transformer modules, composed of attention and feedforward blocks. Finally, a multi layer perceptron enables classification between apoptotic and non-apoptotic sequences.

For the classification of apoptotic sequences, we proposed a Conv-Transformer architecture (Fig. 3). In the proposed architecture, a convolutional module extracts the spatial features of the apoptotic cells, whereas attention-based nlocks evaluates the temporal relationship between consecutive frames.

### 2.3. Training and deployment *in vitro*

As previously described, ADeS is a multiple-block pipeline, and its application and validation to detect apoptotic cells in live-cell imaging follow two main steps: (1) the training of the DL classifier with a target dataset and (2) its deployment on live-cell imaging acquisitions. As opposed to *in vivo* acquisitions, *in vitro* time-lapses are more homogeneous in their content and quality, thus representing the first dataset in order of complexity for the training of ADeS. For this reason, we formulated the learning problem as a binary classification task that assigned non-apoptotic sequences to the class label 0 and apoptotic sequences to the class label 1 (Supplementary Fig. 2A). The class label 0 included instances of healthy nuclei and nuclei undergoing mitotic division (which can resemble apoptotic events).

Successively, to validate the proposed Conv-Transformer architecture for apoptosis classification, we compared it with the performances of a convolutional neural network (CNN), a 3DCNN, and a convolutional long-short term memory (Conv-LSTM) network. To this end, the four models were trained on a dataset containing 13.120 apoptotic and 13.120 non-apoptotic events, using a 0.12 validation split (Table 1). Results show that the frame accuracy of the CNN is low, possibly due to morphological heterogeneity over consecutive frames, and therefore unsuitable for the task. By contrast, the 3DCNN and the Conv-LSTM displayed high sequence accuracy, F1 score and AUC, confirming that the temporal information within frames is pivotal to correctly classifying image sequences containing apoptotic cells. Nonetheless, the proposed Conv-Transformer outperformed both the 3DCNN and the Conv-LSTM, establishing itself as the final DL architecture at the core of ADeS.

**Table 1.** Comparison of deep learning architectures for apoptosis classification. Comparative table reporting accuracy, F1 and AUC metrics for a CNN, 3DCNN, Conv-LSTM, and Conv-Transformer. The classification accuracy is reported for static frames or image-sequences. N.A. stands for non applicable. The last column shows which cell death study employed the same baseline architecture displayed in the table..

| Classifier architecture | Frame accuracy | Sequence accuracy | F1 | AUC | Study |
| --- | --- | --- | --- | --- | --- |
| CNN | 74 % ± 1.3 | N.A. | 0.77 | 0.779 | (Damián La Greca et al. 2021; Verduijn et al. 2021) |
| 3DCNN | N.A. | 91.22 %± 0.15 | 0.91 | 0.924 | - |
| Conv-LSTM | N.A. | 97.42 % ± 0.09 | 0.97 | 0.994 | (Kabir et al. 2022; Mobiny et al. 2020) |
| Conv-Transformer | N.A | 98.27 % ± 0.25 | 0.98 | 0.997 | Our |

Successively, we deployed a preliminary trained network on control movies without apoptotic events to collect false positives that we used to populate the class label 0, thus ensuring a systematic decrease in the misclassification rate (Supplementary Fig. 2B). Using the latter generated dataset, we trained the Conv-Transformer for 100 epochs using an unbalanced training set with a 1:10 ratio of apoptotic to non apoptotic cells (Fig. 4A). After deploying the trained model on 1000 testing samples, the confusion matrix (Fig. 4B) displayed a scant misclassification rate (2.68%), similarly distributed between false positives (1.04%) and false negatives (1.64%). Accordingly, the receiver operating characteristic (ROC) of the model skewed to the left (AUC = 0.99, Fig. 4C). This skew indicates a highly favorable tradeoff between the true positive rate (TPR) and false positive rate (FPR), which the overall predictive accuracy of 97.32% previously suggested (Fig. 4B). Altogether, these metrics suggest an unprecedented accuracy of the DL model in the classification of apoptotic and non-apoptotic sequences. However, they only reflect the theoretical performances of the classifier applied to cropped sequences depicting a single cell at a time.

**Figure 4.**
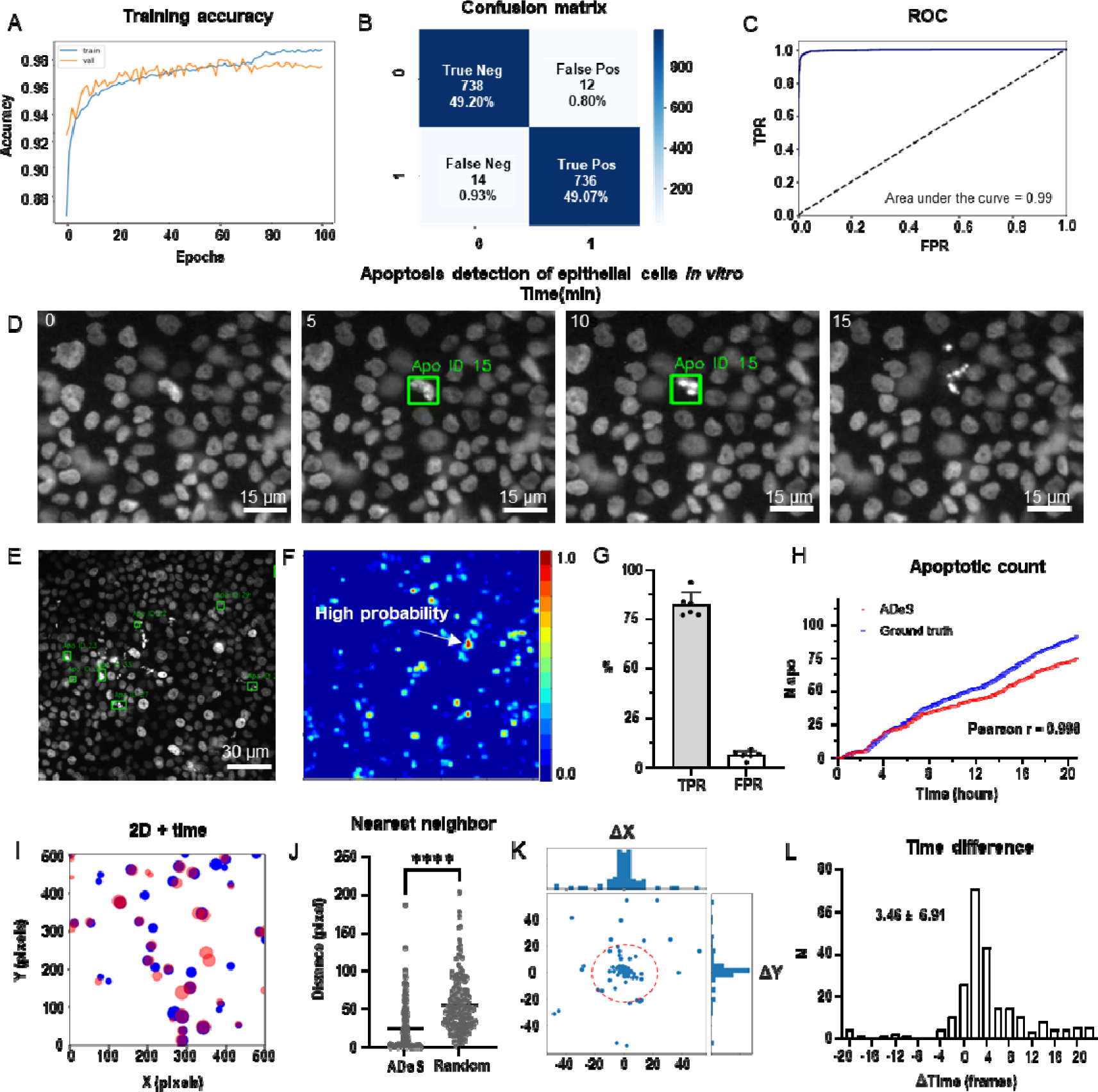
Training and performance in vitro. **A.** Confusion matrix of the trained model at a decision-making threshold of 0.5. **B.** Receiver operating characteristic displaying the false positive rate (specificity) corresponding to each true positive rate (sensitivity). **C**. Training accuracy of the final model after 100 epochs of training. **D.** Representative example of apoptosis detection in a time-lapse acquired in vitro. **E.** Multiple detection of nuclei undergoing apoptosis displays high sensitivity in densely packed field of views. **F.** Heatmap representation depicting all apoptotic events in a movie and the respective probabilities. **G.** Bar plots showing the true positive rate (TPR) and false positive rate (FPR) of ADeS applied to five testing movies, each one depicting an average of 98 apoptosis. **H.** Time course showing the cumulative sum of ground truth apoptosis (blue) and correct predictions (red). **I.** 2D visualization of spatial-temporal coordinates of ground truth (blue) and predicted apoptosis (red). In the 2D representation, the radius of the circles maps the temporal coordinates of the event. **J.** Pixel distance between ADeS predictions and the nearest neighbor (NN) of the ground truth (left) in comparison with the NN distance obtained from a random distribution (right). The plot depicts all predictions of ADeS, including true positives and false positives. **K.** Scatterplot of the spatial distance between ground truth and true positives of ADeS. Ground truth points are centered on the X = 0 and Y = 0 coordinates. **L. Distribution of** the temporal distance (frames) of the correct predictions from the respective ground truth nearest neighbor. Statistical comparison was performed with Mann-Whitney test. Columns and error bars represent the mean and standard deviation respectively. Statistical significance is expressed as: p ≤ 0.05 (*), p ≤ 0.01 (**), p ≤ 0.001 (***), p ≤ 0.0001 (****).

To validate ADeS on full-length microscopy acquisitions, we deployed it on six testing movies that were not part of the training set. Each testing movie had been annotated manually and contained a variable number of ground-truth apoptosis (98 ± 21) and a comparable cell density (1705 ± 124). Moreover, all movies had identical magnification (20x), duration (21 h), and sampling rate (5 min). In order to test ADeS on these movies, we adopted an unbiased approach and we did not hard-tune the hyper-parameters of the model (see Material and Methods), specifying only a stringent confidence threshold (0.995) and a temporal window based on the average duration of the nuclear hallmarks (9 frames). As a result, ADeS could predict the location and timing of the apoptotic nuclei (Fig. 4D, Supplementary Movie 1), enabling the detection of multiple apoptoses in a densely packed field of view (Fig. 4E–F). To quantify these performances, we compared the prediction of ADeS to the annotated ground truths (x,y,t). By doing this, we found that the average TPR, or sensitivity, was 82.01% (ranging from 77% to 92%), while the average FPR was 5.95% (Fig. 4G). The undetected apoptotic events were likely a consequence of the heterogeneity of nuclear fragmentation, which can vastly differ in signal intensity, size, focal plane, and duration (Supplementary Fig.1). Nonetheless, hard tuning the model could further increase the sensitivity without additional training data, such as by adjusting the temporal interval or by lowering the confidence threshold. With respect to the false positives, most were mitotic cells, due to their morphological similarities with aoptotic nuclei. Nevertheless, the FPR was contained, translating into a new false positive every 4 frames (or 20 minutes of acquisition). This rate confirmed that ADeS is overall robust, especially in light of movies depicting 1700 cells per frame.

Concerning the spatial-temporal dynamics, the apoptotic count over time highlighted a tight relationship between ground truth apoptosis and correct detections of ADeS (Fig. 4H). Accordingly, the two curves were divergent but highly correlative (Pearson r = 0.998), proving that ADeS can successfully capture cell death dynamics. A 2D scatterplot (x, y, t = radius; Fig. 4I) visually depicted the spatial-temporal proximity between ADeS and the ground truth, indicating overlap between the two scatter populations. Nearest neighbor (NN) analysis further captureed this relationship; the average distance between all ADeS predictions (true positives + false positives) and the NN in the ground truth was 30 pixels. In contrast, randomly generated predictions had a ground truth NN within a 52-pixel radius (Fig. 4J). Considering instead the true positives only, we observed that they were in close spatial proximity to the ground truth, with most predictions falling within a 20-pixel radius (Fig. 4K). The difference between the predicted timing of apoptosis and the one annotated in the ground truth was also slight, with an average discard of 3.46 frames (Fig. 4L). Interestingly, ADeS showed a bias toward late detections, which is explained considering that operators annotated the beginning of the apoptosis, whereas ADeS learned to detect nuclear disruption, occurring at the end of the process. Altogether, these quantifications indicate that ADeS detects apoptotic nuclei with high spatial and temporal accuracy, establishing a novel comparative baseline for this task.

### 2.4. 3D rotation of the *in vivo* dataset

Upon the successful application of ADeS *in vitro*, the next step in complexity was detecting apoptosis *in vivo* time lapses. The latter is inherently more challenging due to different factors, including high background signal, autofluorescence, and the presence of collagen(Pizzagalli et al. 2018), among others. For this purpose, we re-trained ADeS using the *in vivo* data described in Figure 1. However, one of the main limitations of supervised DL is the need for large datasets, and the finite number of MP-IVM acquisitions and apoptotic instances represented a bottleneck for the training of ADeS. To overcome this limitation, we implemented a custom data augmentation strategy that exploits 3D volumetric rotations, as previously performed in other studies (Xu, Li, and Zhu 2020; Zhuang et al. 2019). Accordingly, each 3D apoptotic sequence underwent multiple spatial rotations and was successively projected in 2D (Fig. 5A). This procedure enabled us to increase the dataset of a 100 fold factor without introducing imaging artifacts, as each volume rotation was a physiological representation of the cell (Fig. 5B).

**Figure 5.**
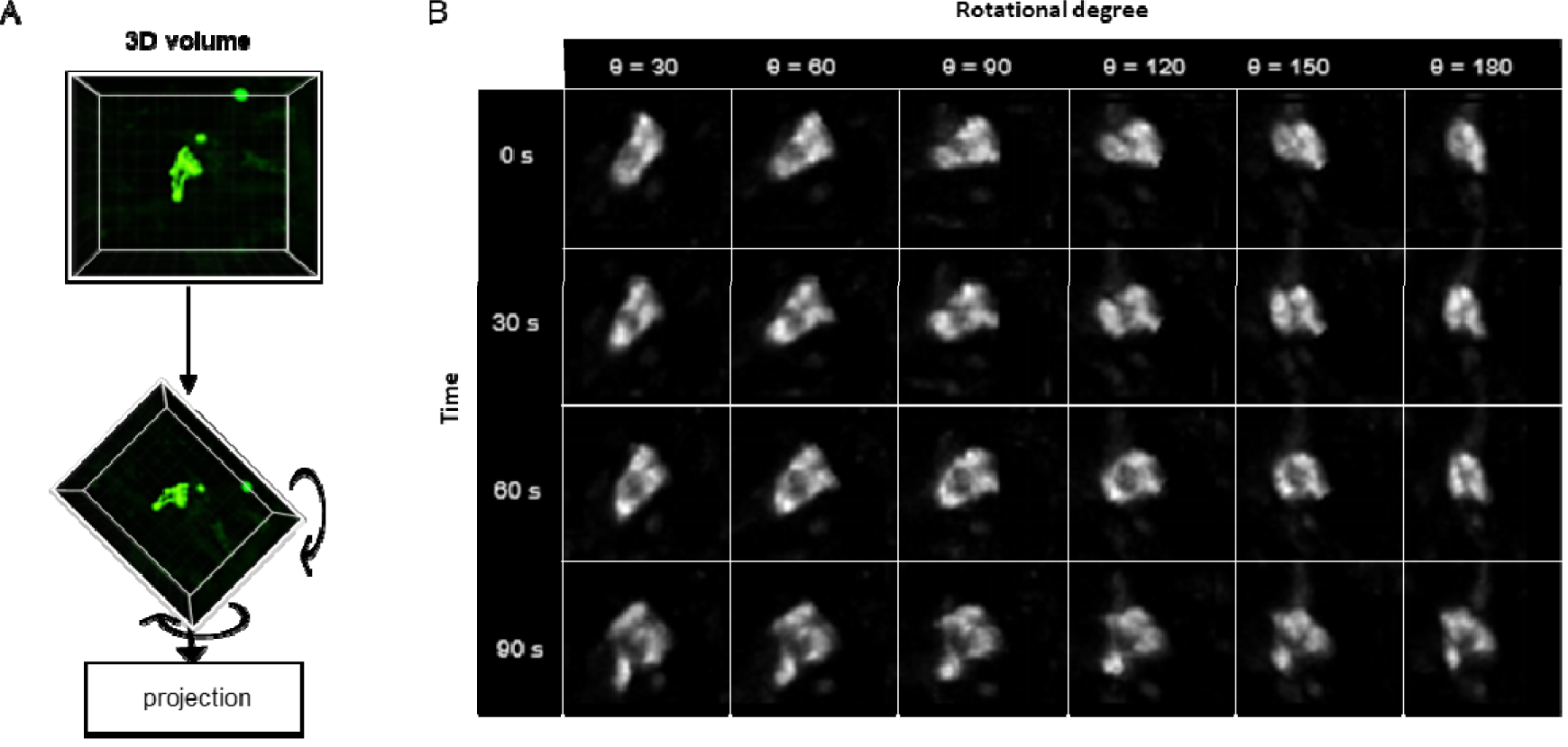
3D rotation of the in vivo dataset. **A** Depiction of a 3D volume cropped around an apoptotic cell. Each collected apoptotic sequence underwent multiple 3D rotation in randomly sampled directions. The rotated 3D images were successively flattened in 2D. **B.** Gallery showing the result of multiple volume rotations applied to the same apoptotic sequence. The vertical axis depicts the sequence over time, whereas the horizontal describes the rotational degree applied to the volumes.

### 2.5. Training and deployment *in vivo*

To train ADeS using the latter rotated *in vivo* dataset, we defined a binary classification task in which ROIs containing apoptotic cells were assigned to the class label 1. In contrast, all remaining ROIs, including healthy cells and background elements, were assigned to the class label 0 (Supplementary Fig. 3A). Subsequently, we trained the DL classifier for 200 epochs. Finally, we performed 5-fold cross-validation according to the ID of the movies (Fig. 6A). The resulting confusion matrix demonstrated a classification accuracy of 97.80% and a 2.20% missclassification rate that is primarily due to type II error (1.80% false negatives) (Fig. 6B). Analogous to the tests *in vitro*, classification *in vivo* proved highly effective in predicting apoptotic and non-apoptotic instances. The ROC of the model, which indicated high sensitivity and a low FPR, supported this favorable result (Fig. 6C).

**Figure 6.**
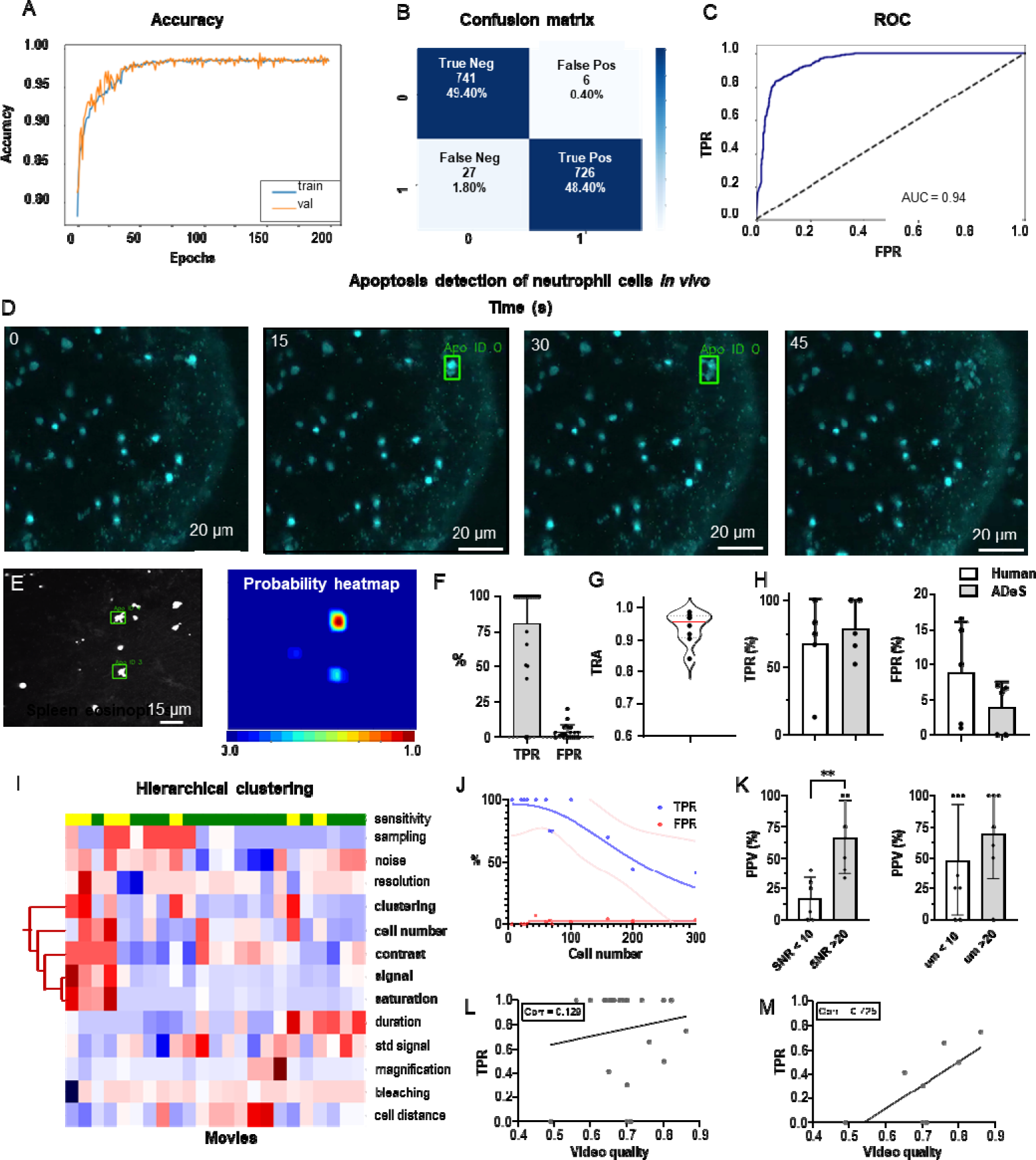
Training and performance in vivo. **A.** Confusion matrix of the trained model at a decision-making threshold of 0.5. **B**. Receiver operating characteristic displaying the false positive rate (FPR) corresponding to each true positive rate (TPR). **C**. Training accuracy of the final model trained for 200 epochs with data augmentations. **D.** Image gallery showing ADeS classification to sequences with different disruption timing. The generated heatmap reaches peak activation (red) at the instant of cell disruption **D.** Representative snapshots of a neutrophil undergoing apoptosis. Green bounding boxes represents ADeS detection at the moment of cell disruption **E.** Representative micrograph depicting the detection of two eosinophil undergoing cell death in the spleen (**left**) and the respective probability heatmap (**right**). **F.** ADeS performances expressed by means of true-positive rate (TPR) and false-positive rate (FPR) over a panel of 23 videos. **G**. TRA measure distribution of the trajectories predicted by ADeS with respect to the annotated ground truth (n = 8) **H**. Comparison between human and ADeS by means of TPR and FPR on a panel of 5 randomly sampled videos **I.** Hierarchical clustering of several video parameters producing two main dendrograms (n = 23). The first dendrogram includes videos with reduced sensitivity and is enriched in several parameters related to cell density and signal intensity. **J.** Graph showing the effect of cell density on the performances expressed in terms of TPR and FPR (n = 13). **K**. Comparison of the positive predictive value between videos with large and small signal to noise ratio (**left**), and videos with large and small shortest cell distance (**right**). **L-M.** Selected video parameters are combined into a quality score that weakly correlates with the TPR in overall data (**M, n = 23**) and strongly correlates with the TPR in selected underperforming data (**N, n = 8**). Statistical comparison was performed with Mann-Whitney test. Columns and error bars represent the mean and standard deviation respectively. Statistical significance is expressed as: p ≤ 0.05 (*), p ≤ 0.01 (**), p ≤ 0.001 (***), p ≤ 0.0001 (****).

We then benchmarked ADeS in the detection task performed on a set of 23 MP-IVM acquisitions of immune cells undergoing apoptosis. Unlike *in vitro* settings, *in vivo* acquisitions displayed high variability in cell number, auto-fluorescence, signal intensity, and noise levels (Supplementary Fig. 3B). Still, ADeS correctly predicted the location and timing of cells undergoing apoptosis (Fig. 6H, Supplementary Movie 2), indicating its robustness to increasingly populated fields of view (Supplementary Fig. 3C). In addition, we successfully applied the pipeline to neutrophils imaged in the lymph node (Fig. 6D) and eosinophils in the spleen (Fig. 6E). By comparing ADeS predictions with the annotated ground truths, we found that our pipeline detected apoptotic events with a TPR of 81.3% and an FPR of 3.65% (Fig. 6F). The detections, provided in the form of bounding boxes and trajectories, indicated the coordinates and duration of the events. Hence, to measure how close they were to the annotated trajectories, we employed the tracking accuracy metric (TRA), a compound measure that evaluates the similarities between predicted and ground truth trajectories. The average TRA was above 0.9, indicating the high fidelity of the trajectories predicted by ADeS (Fig. 6G).

Next, we compared ADeS to human annotation performed by three operators on five testing movies. As a result, ADeS displayed an upward trend of the TPR and a downward trend of the FPR. However, we found no significant difference in the TPR and FPR (Fig. 6H). Regardless, ADeS performances appeared to be distributed across two distinct groups: a predominant group with an average sensitivity of 100% ( > 75% range) and a smaller group with an average sensitivity of 53% (41-75% range, Fig. 6H). To understand this discrepancy, we applied hierarchical clustering to the testing videos according to their imaging properties and biological content (Fig. 6I), thus generating two major dendrograms. The first dendrogram mostly contained videos with reduced sensitivity (yellow) and was defined by a high cell number, high noise levels, short cell distance, and a saturated and fluctuating image signal. Most notably, the cell number played a crucial role in overall performance, as reflected in the fact that an increment of this parameter resulted in a pronounced decrease in the TPR and a moderate increase in the FPR (Fig. 6J). Incidentally, the positive predictive value (PPV) was significantly lower in videos with poor signal-to-noise ratio (SNR) and, although not statistically significant, the PPV was lower when the signal standard deviation was higher (Fig. 6K, Supplementary Movie 3). As similar findings were observed *in vitro* (Supplementary Fig. 4), we hypothesized that the quality of a movie predicts ADeS performance. Hence, we combined the parameters highlighted by the clustering analysis (Fig. 6I) into a single score ranging from zero to one (one indicating the highest and ideal score) and, in doing so, found there to be a weak correlation between the video quality and the sensitivity of ADeS (Fig. 6L). However, this trend was evident only when we considered videos with suboptimal sensitivity; indeed, in these cases, we found a strong correlation (0.72), confirming that the video quality partially explains the observed performances (Fig. 6M).

Finally, we evaluated how the biological variability *in vivo* could affect the readout of ADeS, defining nine distinct biological categories, including apoptotic cells, healthy cells, and background elements. For all biological categories, the classification accuracy was above 80%, except for overlapping cells and cells with high membrane plasticity (Supplementary Fig. 3D).

### 2.6. Comparison with the state-of-the-art

To compare the performance of ADeS with other state-of-the-art algorithms for cell death quantification, we conducted a comprehensive literature review. For each study, we reported the attained classification accuracy, the experimental setup, the architecture of the classifier, the capability of detecting cell death events in movies, and the number of cell deaths in the training set (Table 2). Initial results indicate that ADeS achieved the highest classification accuracy, but a direct comparison in terms of accuracy is not meaningful due to the differences in datasets, including distinct cell types, different types of cell death, and varying dataset sizes. For a more appropriate benchmark, we refer to Table 1, which shows that our classifier outperformed the baseline re-implementations of the main classifiers used in other studies.

**Table 2.** Comparison of cell death identification studies. Table reporting all studies on cell death classification based on machine learning. For each study, we included the reported classification accuracy, the experimental conditions of the studies, the target input of the classifier, and the capability of performing detection on static frames or microscopy time-lapses. Met conditions are indicated with a green check. Moreover, for each study we reported the architecture of the classifier and the number of apoptotic cells in the training set. N.A. stands for not available and indicates that the information is not reported in the study.

| Study | Input of the classifier | Reported classification accuracy | *In vitro* | *In vivo* | Detection In frame | Detection in movies | Classifier architecture | N cell death |
| --- | --- | --- | --- | --- | --- | --- | --- | --- |
| Our | Frame sequence | 98.27 % | ? | ? | ? | ? | Conv-Transformer | 13'120 |
| (Jin et al. 2022) | Frame | 93 % | ? | ? | ? | ? | Logistic regression | N.A. |
| (Verduijn et al. 2021) | Frame | 87 % | ? | ? | ? | ? | VGG-19 | 19'339 |
| (Kabir et al. 2022) | Frame sequence | 93 % | ? | ? | ? | ? | ResNet101-LSTM | 3'172 |
| (Damián La Greca et al. 2021) | Frame | 96.58 % | ? | ? | ? | ? | ResNet50 | 11'036 |
| (Mobiny et al. 2020) | Frame sequence | 93.8 % | ? | ? | ? | ? | CapsNet-LSTM | 41'000 |
| (Kranich et al. 2020) | Frame | 93.2 % | ? | ? | ? | ? | CAE-RandomForest | 27'224 |
| (Vicar et al. 2020) | Frame sequence | N.A. | ? | ? | ? | ? | biLSTM | 1'745 |
| (Jimenez-Carretero et al. 2018) | Frame | N.A. | ? | ? | ? | ? | R-CNN | 255'215 |

From Table 2, we observe that ADeS is the only algorithm for cell death quantification that has been applied *in vivo*. Additionally, only ADeS and the study by Vicar and colleagues(Vicar et al. 2020) effectively detected apoptotic cells in fully uncropped microscopy movies, which is a significant achievement given the computational challenge associated with the task. However, Vicary and colleagues relied on the temporal analysis of cell trajectories, while ADeS used vision-based methods to directly analyze consecutive frames of a movie. As a result, ADeS offers a comprehensive and pioneering pipeline for effectively applying vision-based classifiers to detect cell death in imaging time-lapses.

### 2.7. Applications for toxicity assay *in vitro*

A common application of cell death staining is the evaluation of the toxicity associated with different compounds(Atale et al. 2014; Schmid, Uittenbogaart, and Jamieson 2007) or the efficacy of an apoptotic-inducing treatment. Here, we show that ADeS has analogous purposes and can effectively quantify the toxicity of different compounds *in vitro*. For this application, we grew epithelial cells *in vitro*, treating them with PBS and three increasing concentrations of doxorubicin, a chemotherapeutic drug that elicits apoptosis in the epithelium(Eom et al. 2005). Epithelial cells were seeded with the same density of cells per well, and all four conditions had the same confluence before the treatment. However, at 24 h. post-acquisition, the number of survivor cells was inversely proportional to the doxorubicin concentration (Fig. 7A). We confirmed this trend using ADeS (Supplementary Movies 4–7), which measured the lowest mortality after 24 h. in PBS (62 cells), followed by doxorubicin concentrations of 1.25 μM (95 cells), 2.50 μM (167 cells), and 5.00 μM (289 cells). Moreover, ADeS predicted distinct pharmacodynamics (Fig. 7B), which can define the drug concentration and experimental duration required to reach a specific effect in the apoptotic count. To this end, each time point in Figure 7B also defines a dose-response relationship. Here we provide two dose-responses curves at 5 h. and 24 h. post-treatment, showing different pharmacodynamics (EC50 5h = 2.35, Hill slope 5h = 3.81, EC50 24h = 4.47, Hill slope 24h = 1.93, Fig. 7C–D). Notably, the fit can project the dose-responses for higher drug concentrations, predicting the maximum effect size at a given time. For instance, at 24 h. post treatment, a 10 μM titration attains 86% of the maximum effect (456 apoptotic cells), whereas a further increase in the concentration of the drug leads only to a moderate increase of the toxicity (Fig. 7E). We argue that this approach helps to maximize the effect of a drug on a designated target, while minimizing collateral damage done to non-target cells. For instance, in chemotherapies employing doxorubicin, apoptosis of epithelial cells is an undesired effect. Therefore, researchers can select a titration of the drug and a duration of the treatment that does not affect the epithelium yet still positively affects the tumor. Finally, we also demonstrated the reproducibility of the toxicity assay by targeting another cell type (T-cells) treated with a different apoptotic inducer (staurosporine, Supplementary Fig. 5).

**Figure 7.**
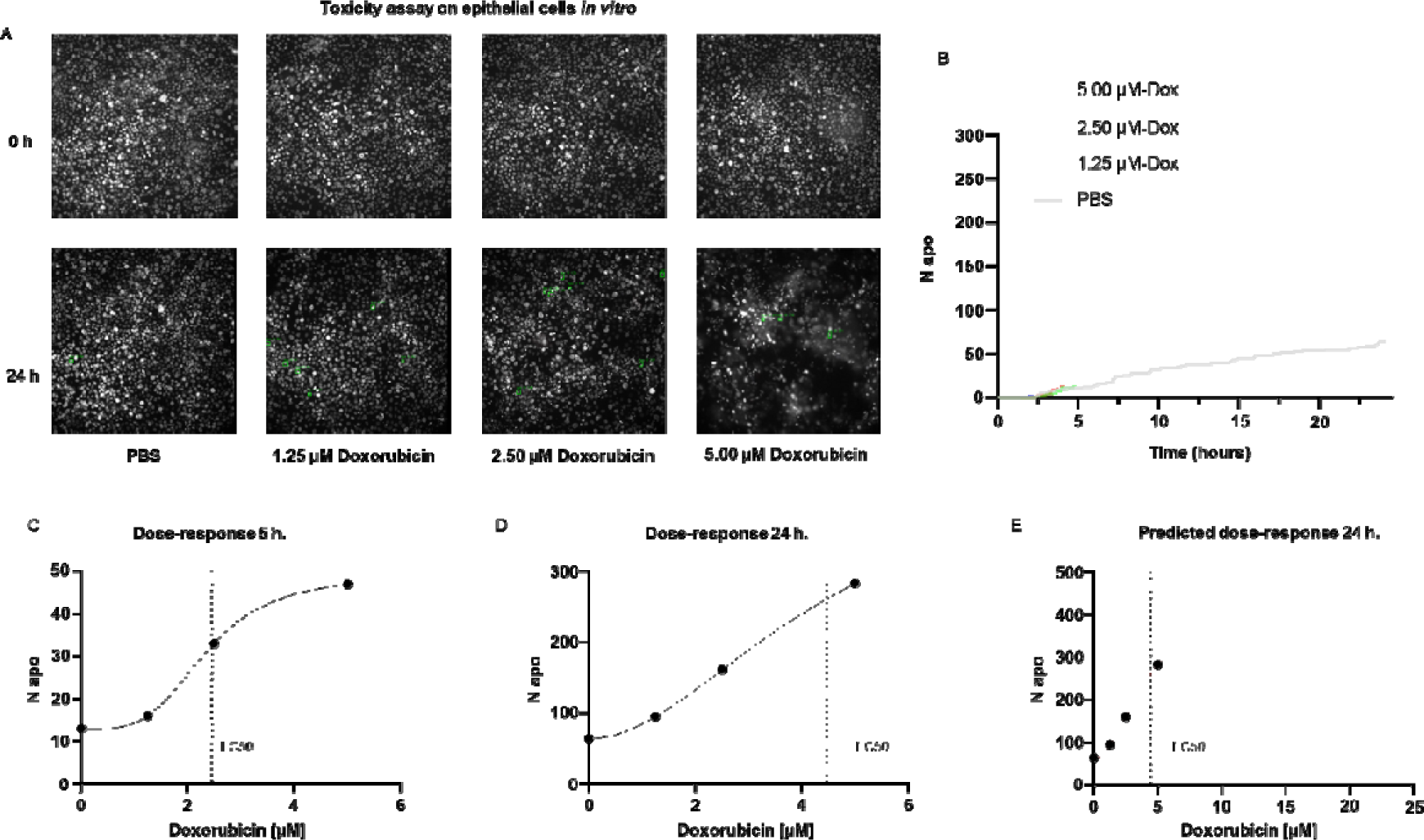
Applications for toxicity assay in vitro. **A.** Representative snapshots depicting epithelial cells in vitro at 0 and 24 hours after the addition of PBS and three increasing doses of doxorubicin, a chemotherapeutic drug and apoptotic inducer **B**. Plot showing the number of apoptotic cells detected by ADeS over time for each experimental condition. **C-D**. Dose-response curves generated from the drug concentrations and the respective apoptotic counts at 5 h. and 24 h.post-treatment. Vertical dashed lines indicates the EC50 concentration. **E.** Dose-response curve projected from the fit obtained in (**D**). The predicted curve allows to estimate the response at higher drug concentrations than the tested ones.

### 2.8. Measurement of tissue dynamics *in vivo*

To test the application of ADeS in an *in vivo* setting, we applied it to study the response of bystander cells following apoptotic events in the lymph nodes of mice treated with an influenza vaccine. We computed the spatial and temporal coordinates of a neutrophil undergoing apoptosis (Fig. 8A), which, combined with the tracks of neighboring cells, allowed us to characterize cellular response patterns following the apoptotic event. Among other parameters, we observed a sharp decrease in the distance between the neighboring cells and the apoptotic centroid (Fig. 8B) in addition to a pronounced increase in the instantaneous speed of the cells (Fig. 8C).

**Figure 8.**
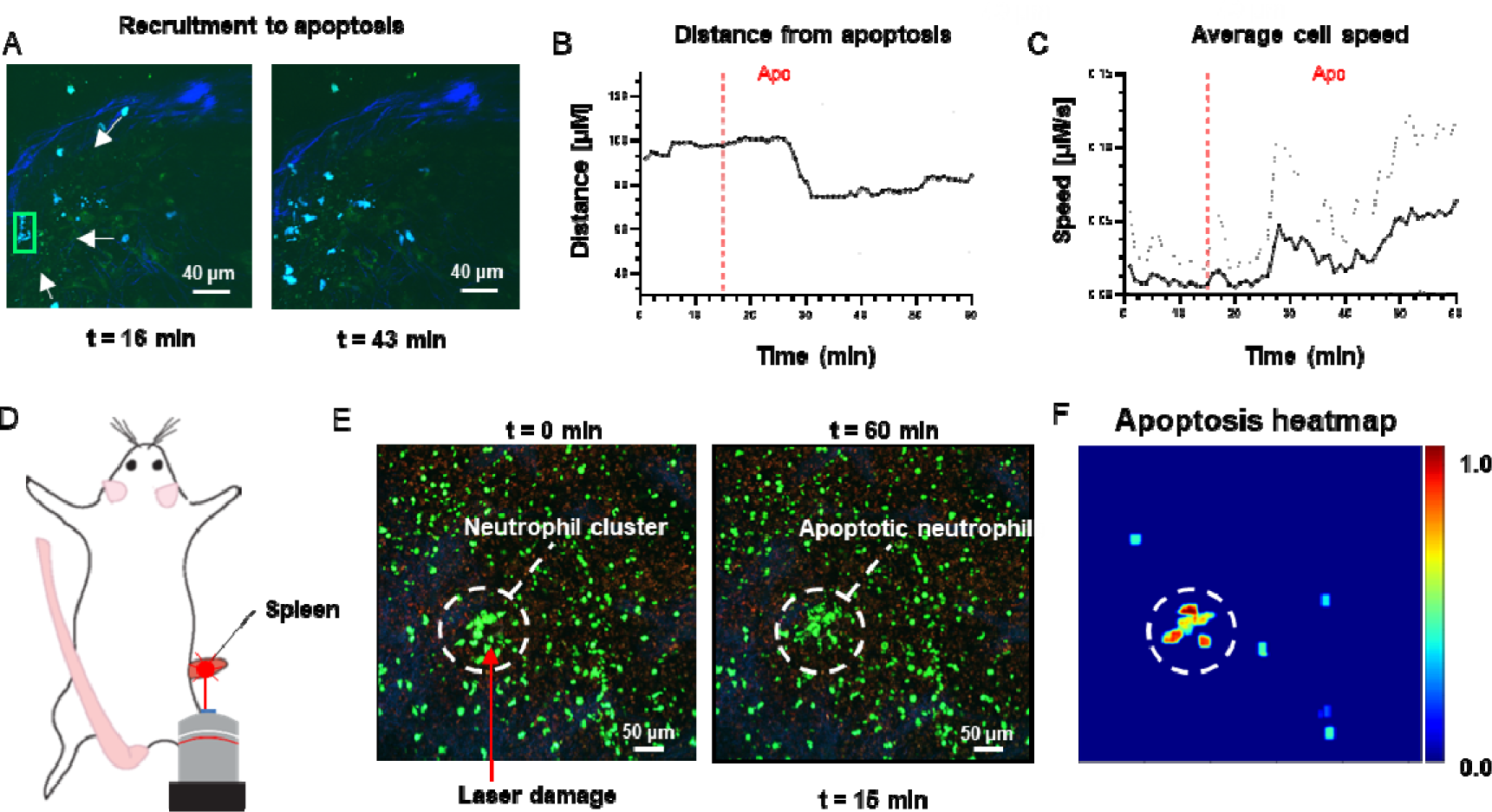
Measurement of tissue dynamics in vivo. **A.** Intravital 2-photon micrographs showing ADeS detection of an apoptotic neutrophil (Blue, left) and the subsequent recruitment of neighboring cells (right) in the popliteal LN at 19 h. following influenza vaccination. **B**. Plot showing the distance of recruited neutrophils with respect to the apoptotic coordinates over time (n = 22) **C.** Plot showing the instantaneous speed of recruited neutrophils over time (n = 22). The dashed vertical lines indicate the instant in which the apoptotic event occurs. Gray area defines the boundaries of maximum and minimum values. **D**. Schematic drawing showing the intravital surgical set up of a murine spleen after inducing a local laser ablation. **E**. Intravital 2-photon micrographs showing the recruitment of GFP-expressing neutrophils (Green) and the formation of a neutrophil cluster (red arrows) at 60 min after photo burning induction. **F**. Application of ADeS to the generation of a spatiotemporal heatmap indicating the probability of encountering apoptotic events in the region affected by the laser damage. The dashed circle indicates a hot spot of apoptotic events.

Successively, we evaluated the detection of apoptotic cells following laser ablation in the spleen of an anesthetized mouse (Fig. 8D). Previous research has employed this method to study immune cell responses to tissue damage(Uderhardt et al. 2019). The insult caused prompt recruitment of neutrophils, leading to the formation of a local swarm (Fig. 8E, left). After that, the neutrophils within the swarm underwent apoptotic body formation in a coordinated manner (Fig. 8E, right). To quantify this event, we processed the generated time-lapse with ADeS, resulting in a probability map of apoptotic events throughout the acquisition (x,y,t,p; Fig. 8F). Accordingly, the location with the highest probability corresponded to the area damaged by the laser, while the visual representation of the probability map enabled us to infer the morphology and location of the swarm. This result demonstrates the potential application of ADeS in digital pathology, showing how the distribution of apoptotic events throughout the tissue can identify areas enriched by cell death events.

## 3. Discussion

Automated bio-image analysis obviates the need for manual annotation and avoids bias introduced by the researcher. In this regard, recent studies showed the promising usage of DL to classify static images(Jimenez-Carretero et al. 2018; Kranich et al. 2020; Verduijn et al. 2021) or time-lapses containing single apoptotic cells(Mobiny et al. 2020). However, these approaches are unsuitable for microscopy time-lapses because they do not address two fundamental questions: the location, over the whole field of view, at which an event occurs, and its duration. These questions define a detection task(Zhao et al. 2019) in space and time, which has a computational cost that can rapidly grow with the size and lenght of a movie. Moreover, live-cell imaging data present specific challenges which further increase the difficulty of detection routines, including densely packed fields of view, autofluorescence, and imaging artifacts(Pizzagalli et al. 2018).

Consequently, computational tools to effectively detect apoptotic events in live-cell imaging remained unavailable. Thus, we created an apoptosis detection pipeline that could address the abovementioned challenges *in vitro* and *in vivo*. In this regard, ADeS represents a crucial bridge between AR and bioimaging analysis, being the first apoptosis detection routine with demonstrated applicability to full microcopy time-lapses. In addition, we presented two comprehensive and curated datasets encompassing multiple cell types, fluorescent labels, and imaging techniques to encourage reproducibility and foster the development of apoptosis detection routines.

In human activity recognition benchmark, 3DCNNs(Vrskova et al. 2022), two-streams networks(Ye et al. 2019), and RNNs(Mohd Noor, Tan, and Ab Wahab 2022) have proved to score the highest accuracy on most kinetic datasets(Ullah et al. 2021). However, in most studies for the classification of apoptosis, authors unanimely employed RNNs such as Conv-LSTMs. This choice, although produced valid results, is not necessarily optimal for the task. In this regard, Ullah and colleagues highlighted that the performances of different DL architectures are highly dependent on the AR dataset(Ullah et al. 2021). Therefore, selecting the most suitable one is only possible after an extensive benchmark. In our comparison, we demonstrated for the first time that attention-based networks are suitable for the classification and detection of apoptotic events. Accordingly, our Conv-Transformer network outperformed DL architectures previously employed in other studies, including 3DCNNs and RNNs. This result established a landmark in the application of attention-based networks in AR for live-cell imaging. Moreover, it suggests the possible benefits of employing transformers for the classification of different biological activities other than cell death.

Similar to most diagnostic tools, ADeS displayed a tradeoff between sensitivity (TPR) and specificity (1 - FPR), which is a known challenge in binary classification(Pang et al. 2021). This tradeoff can be attributed to the fact that apoptosis is rare in normal physiological conditions, leading to a high degree of class imbalance during training. As a result, the choice of the training set had a significant impact on the performances of ADeS. For instance, we highlighted the importance of a training and validation set that included challenges related to real live-cell imaging acquisitions, such as overlapping cells and low signal-to-noise samples. Including these challenges instances enabled ADeS to attain low misclassification rate and robust real-life performanes. Nonetheless, we observed residual misclassifications due to shared similarities between healthy and apoptotic cells. For instance, *in vitro* mitotic divisions could mislead the detection of apoptotic nuclei, while *in vivo*, overlapping cells were sometimes mistaken for apoptotic cells. Therefore, to effectively address these challenges, it is crucial to implement strategies to increase the representativeness of the dataset, such as integrating multiple data sources and data augmentation techniques.

From a biological perspective, ADeS has multiple applications in fundamental and clinical research. Among other advantages, it can provide insights into pivotal cell death mechanisms, monitor the therapies used to modulate apoptosis in various diseases and characterize the toxicity of different compounds. In this regard, ADeS readout is analogous to standard fluorescent probes for apoptosis detection, with the advantage that it can be applied directly to nuclear or cytoplasmic staining without the need of additional fluorescent reporters. Therefore, ADeS avoids using any additional acquisition channel, which can be used for multiplexing purposes. Moreover, common probes(Atale et al. 2014; Kyrylkova et al. 2012; Loo 2011; Sun et al. 2008; Vermes et al. 1995) flag early apoptosis stages, activated up to several minutes before the point at which morphological changes in the cell(Green 2005; Takemoto et al. 2003); meanwhile, these cells can reverse the apoptotic process (Geske et al. 2001; Masri and Chandrashekhar 2008; H. L. Tang et al. 2009). By contrast, ADeS indicates the exact instant of cell disruption, thus adding specificity to the spatial-temporal dimension. For these reasons, we suggest that ADeS can complement the information provided by classic apoptotic biomarkers, which will prove advantageous in experimental assays where the temporal resolution delivers more information than the sole apoptotic count. Moreover, ADeS can be usefully applied in processing high-throughput live-cell imaging, minimizing annotation time and research bias.

Finally, in tissue dynamics the spatial-temporal activity of cells can reveal connections between signaling pathways and the fate decision of individual cells, such as mitosis or apoptosis(Gagliardi et al. 2021). These intricate systems can display complex dynamics, which can be better comprehended incorporating spatial and temporal coordinates provided by ADeS. Consequently, we propose that integrating these spatial-temporal characteristics with experimental observations could lay the groundwork for understanding the mechanism governing complex signaling pathways. Furthermore, we contend that this information has the potential to facilitate the development of predictive models, establishing a correlation between specific cell death dynamics and the underlying stimuli. This, in turn, could serve as the foundation for innovative diagnostic tools capable of inferring the cause of cell death.

(Fesik 2005; Hotchkiss and Nicholson 2006)( *15*, *16*)[15], [16] ^15,16^In conclusion, ADeS constitutes a novel solution for apoptosis detection that combines state-of-the-art microscopy and DL. Its successful implementation represents a step towards the general application of AR methods to live-cell imaging. By bridging these two distinct fields, ADeS leverages succesfully the benefits of automated routines. Further work could expand the proposed pipeline to encompass diverse cell populations, various types of cell death, and potentially broader cellular activities.

## 4. Material and methods

### 4.1. MCF10A cell line and image acquisition

The normal-like mammary epithelial MCF10A cells (provided by Joan Brugge(Debnath, Muthuswamy, and Brugge 2003)), stably expressing the nuclear marker, were generated as previously described(Gagliardi et al. 2021). Briefly, the nuclear marker H2B-miRFP703, provided by Vladislav Verkhusha (Addgene plasmid # 80001)(Shcherbakova et al. 2016), was subcloned in the PiggyBac plasmid pPBbSr2-MCS. After cotransfection with the transposase plasmid(Yusa et al. 2011), cells were selected with 5 µg/ml Blasticidin and subcloned. For time-lapse imaging, the cells were seeded on 5 µg/ml fibronectin (PanReac AppliChem) coated 1.5 glass-bottom 24 well plates (Cellvis) at 1 x 10^5^ cells/well density. After 48 hours, when the optical density was reached, the confluent cell monolayer was acquired every 1 or 5 minutes for several hours with a Nikon Eclipse Ti inverted epifluorescence microscope with 640nm LED light source, ET705/72m emission filter and a Plan Apo air 203 (NA 0.8) or a Plan Apo air 403 (NA 0.9) objectives. The collection of biological experiments used in this study includes different stimulation of apoptosis, such as growth factors, serum starvation and doxorubicin at various concentrations.

### 4.2. Apoptosis induction of MCF10A cells with doxorubicin

Normal-like mammary epithelial MCF10A cells were grown in 24 well glass coated with fibronectin with a seeding of 1×10^5^ cells/well. After two days, cells were starved for three hours and treated with doxorubicin at 1.25, 2.50, and 5.00 μM concentrations.

### 4.3. Mice

Prior to imaging, mice were anesthetized with a cocktail of Ketamine (100 mg/Kg) and Xylazine (10 mg/Kg) as previously described(Sumen et al. 2004). All animals were maintained in specific pathogen-free facilities at the Institute for Research in Biomedicine (Bellinzona, CH). All the experiments were performed according to the regulations of the local authorities and approved by the Swiss Federal Veterinary Office.

### 4.6. **Intravital Two-Photon Microscopy.**

Surgery in the popliteal lymph node was performed as previously reported(Miller et al. 2004). The exposed organs were imaged on a custom up-right two-photon microscope (TrimScope, LaVision BioTec). Probe excitation and tissue second-harmonic generation (SHG) were achieved with two Ti:sapphire lasers (Chamaleon Ultra I, Chamaleon Ultra II, Coherent) and an optical oscillator that emits in the 1,010–1,340 nm range (Chamaleon Compact OPO, Coherent) and has an output wavelength between 690–1,080 nm.

### 4.5. Neutrophil isolation from mouse bone marrow

Bone marrow samples were extracted via flushing with PBS from the long bones of UBC-GFP mice (https://www.jax.org/strain/004353). Then, the bone marrow was filtered through a 40um strainer and resuspended in PBS. Primary bone marrow neutrophils were isolated with Ficoll gradient and resuspended in PBS.

### 4.6. T-cell culture in a 3D collagen matrix

Human CD4+ T cells were isolated from the PBMC fraction of healthy donors obtained from NetCAD (Canadian Blood Services). Cell purity was above 95%. Naïve CD4+ T cells were activated by adding Dynabeads coated with anti-human CD3e/CD28 antibody (1:1 bead:cell ratio, Life Technologies Cat #11131D) in RPMI1640 supplemented with 10% FBS (VWR Seradigm Cat #1500-500), 2 mM GlutaMAX (Gibco Cat #3050-061), 1mM sodium pyruvate (Corning Cat #25-000-CI) and 10mM HEPES (Sigma-Aldrich Cat #H4034). After two days, beads were removed and cells were cultured for another 4-6 days in a medium containing 50 IU/mL human rIL-2 (Biotechne Cat #202-IL-500), keeping cell density at 2 x 10^5^ cells/mL. Cells were used for all experiments between days 6 to 8. All work with human blood has been approved by the University of Manitoba Biomedical Research Ethics Board (BREB).

### 4.7. Apoptosis live-cell imaging of T-cells in 3D collagen chambers

T cells were labeled at day 6-8 using CMAC (10µM) cell tracker dye (Invitrogen) and glass slide chambers were constructed as previously described(Lopez et al. 2019, 2022). Briefly, 2 x 10^6^ cells were mixed in 270µL of bovine collagen (Advanced Biomatrix cat #5005-100ML) at a final concentration of 1.7 mg/mL. Collagen chambers were solidified for 45 minutes at 37°C / 5% CO2 and placed onto a custom-made heating platform attached to a temperature control apparatus (Werner Instruments). For the induction of apoptosis, 1µM of Staurosporine (Sigma Cat #569397-100UG) and 800ng of TNF-a (Biolegend Cat #570104) in 100µL RPMI were added on top of the solidified collagen. Cells were imaged as soon as the addition of apoptosis inducers using a multiphoton microscope with a Ti:sapphire laser (Coherent), tuned to 800 nm for optimized excitation of CMAC. Stacks of 13 optical sections (512 x 512 pixels) with 4 mm z-spacing were acquired every 15 seconds to provide imaging volumes of 44mm in depth (with a total time of 60 - 120 minutes). Emitted light was detected through 460 / 50nm, 525 / 70 nm, and 595 / 50 nm dichroic filters with non-descanned detectors. All images were acquired using the 20 x 1.0 N.A. Olympus objective lens (XLUMPLFLN; 2.0mm WD).

### 4.8. Data Processing and Image Analysis

The raw video data, composed by uint8 or uint16 TIFFs, were stored as HDF5 files. No video pre-processing was applied to the raw data before image analysis. Cell detection, tracking, and volumetric reconstruction of microscopy videos were performed using Imaris (Oxford Instruments, v9.7.2). The resulting data were further analyzed with custom Matlab and Python scripts (see code availability section).

### 4.9. Apoptosis annotation of epithelial MCf10A cells *in vitro*

We manually annotated apoptotic events of MCF10A cells by visual inspection of the movies. The annotation was done by observing the morphological changes associated with apoptosis (e.g. nuclear shrinkage, chromatin condensation, epithelial extrusion, nuclear fragmentation) across multiple consecutive frames. Using a custom Fiji(Schindelin et al. 2012) macro, we automatically stored x and y centroids of the apoptotic nucleus. The time t of each apoptotic annotation was defined as the beginning of nuclear shrinkage.

### 4.10. Generation of the *in vitro* training dataset

The 16-bit raw movies were min-max scaled to the 0.001 and 0.999 quantiles and downsampled to 8-bit resolution. Using the database of manually labeled coordinates of apoptotic events (x,y,t), we extracted crops with 59 x 59 pixels resolution (2x scaling for the field of view acquired with the 20x objective). Seven time-steps of the same location were extracted, with linear spacing from −10 minutes to +50 minutes relative to the apoptosis annotation. This time frame was chosen to capture the cell before the onset of apoptosis, and the morphological changes associated with apoptosis (nuclear shrinkage, decay into apoptotic bodies, extrusion from epithelium). The resulting image cube has dimensions of 59 x 59 x 7. To create the training data for the non-apoptotic class, we excluded areas with an annotated apoptotic event with a safety margin from the movies. From the remaining regions without apoptoses, we extracted image cubes from cells detected with StarDist(Jacquemet et al. 2020) and from random locations. The random crops also included debris, apoptotic bodies from earlier apoptotic events, empty regions, and out-of-focus nuclei.

### 4.11. Apoptosis annotation of leukocyte cells *in vivo*

Three operators independently annotated the videos based on selected morphological criteria. To label apoptotic cells, the annotators considered only the sequences of cells that displayed membrane blebbing followed by apoptotic bodies formation and cell disruption (Fig. 2B). For each frame in the apoptotic sequence, the operators placed a centroid at the center of the cell with the Imaris “Spots” function, generating an apoptotic track. Successively, ground truth tracks were generated according to a majority voting system, and 3D volume reconstruction was performed on ground truth cells using the Imaris “Surface” function. Nearby non-apoptotic cells were also tracked. In addition, other non-apoptotic events were automatically sub-sampled from regions without apoptotic cells.

### 4.12. 3D rotation of the *in vivo* annotations

*In vivo* annotations presented a class unbalance in favor of non-apoptotic cells, with a relative few apoptotic instances. Hence, to compensate for this bias, we produced several representations of the raw data by interpolating the raw image stacks in 3D volumes and rotating them in randomly sampled directions, with rotational degrees between 0° and 45°. After each manipulation, the rotated volume underwent flattening by maximum projection and symmetric padding to preserve the original dimension. The 2D images were successively resized and cropped to match the 59 x 59 pixels input of the classifier. Finally, the training sequences were saved as uint8 gray-scale TIFFs files.

### 4.13. Generation of the *in vitro* and *in vivo* training datasets

To detect apoptotic cells in microscopy acquisitions, we defined a 2D binary classification task in which apoptotic events are labeled with class 1, while non-apoptotic events belonged to the class label 0. The resulting unprocessed data consisted of frame sequences composed of 3D crops. The content of the class label 0 in vitro included: healthy nuclei, background, cell debris and mitotic cells. The content of the class label 0 *in vivo* included: motile cells, arrested cells, highly deformed cells, overlapping cells, cell debris or blebs, empty background, noisy background, and collagen.

### 4.14. Data augmentation and data loader

Given the varying length of the training sequences contained in the TIFFs, upon training, we used a custom data loader that uniformly samples the input data and produces sequences with a fixed number of frames. The fixed number of frames was set to 5, corresponding to the frame-length of the shortest apoptotic sequence. During training, each sample underwent horizontal shift, vertical shift, zoom magnification, rotation, and flipping. All data augmentations were performed in python using the Keras library.

### 4.15. Deep learning architecture

As a deep learning classifier, we employed a custom architecture relying on time-distributed convolutional layers stacked on top of a transformer module (Conv-Transformer). The input size consists of 5 single-channel images with 59×59 pixel size. The convolutional network has three layers of size 64, 128, and 256 length. Each layer has a 3×3 kernel, followed by Relu activation, batch normalization, and a dropout set to 0.3. The inclusion of padding preserves the dimension of the input, while 2D max-pooling is at the end of each convolutional block. After 2D max pooling, the output is passed to a transformer module counting 6 attention heads, and successively to a fully connected decision layer. The fully connected network has four layers with 1024, 512,128, and 64 nodes, each one followed by Relu activation and a 0.3 dropout layer. The last layer is a softmax activation, which predicts a decision between the two classes.

### 4.16. Training and hyper-parameters

Our model was trained in TensorFlow with Adam optimizer, using binary cross-entropy loss and an initial learning rate of 0.0001. The optimal mini-batch size was 32, and the number of training epochs was 200. In training mode, we set a checkpoint to save the model with the best accuracy on the validation dataset, and a checkpoint for early stopping with patience set to 15 epochs. In addition, the learning rate decreased when attending a plateau.

### 4.17. ADeS deployment

For the deployment of the classifier on microscopy videos, we generative region proposals using the selective search algorithm, obtaining a set of ROIs for each candidate frame of the input movie. For each ROI computed by the region proposal at time t, a temporal sequence is cropped around t and classified with the Conv-Transformer. The resulting bounding boxes are filtered according to a probability threshold and processed with the non-maxima suppression utils from Pytorch. Consecutive bounding boxes classified as apoptotic are connected using a custom multi-object tracking algorithm based on Euclidean distance. The generated trajectories are filtered by discarding those with less than two objects.

### 4.18. Default and user-defined parameters

ROIs detected with the region proposal are filtered according to their size, discarding the ones with edges below 20 pixels and above 40 pixels. Furthermore, a threshold on intensity is applied to exclude uint8 patches with an average brightness below 40. Upon classification, a temporal window corresponding to the expected duration of the apoptotic event is set by the user (9 frames by default). This temporal window is subsampled to match the number of input frame of the classifier (5). The filtering of the predictions depends on a user-specified threshold, which by default corresponds to 0.95 *in vivo* and 0.995 *in vitro*. Non-maxima suppression is based on the overlapping area between bounding boxes, set to 0.1 by default. The centroid tracking has the following adjustable parameters: gap and distance threshold. The “gap” parameter, set to three frames, specifies for how long a centroid can disappear without being attributed a new ID upon reappearance. A threshold on the distance, set by default to 10 pixels, allows the connection of centroids within the specified radius. All the reported quantifications had default parameters.

### 4.19. Statistical analyses

Statistical comparisons and plotting were performed using GraphPad Prism 8 (Graphpad, La Jolla, USA). All statistical tests were performed using non-parametric Kruskal-Wallis test or Mann-Witney test For significance, p value is represented as * when p < 0.05, ** when p < 0.005 and *** when p < 0.0005.

## 5. Data availability

Imaging data: https://zenodo.org/uploads/10260643

Code: https://github.com/mariaclaudianicolai/ADeS

## 5. Aknowledgements

We would like to thank Dr. Coral Garcia (IQS, Barcelona, Spain) for the support in generating graphical content. Moreover, we would like to aknowledge Gabriele Abbate (IDSIA, Lugano, Switzerland) for his help during an early implementation of the DL classifier.

## 5.2. Authors contributions

Conceptualization: SG, DU, AP

Methodology: AP, DU

Experiments: PG, PA, MP, RZ, PL

Data annotation: PG, AP, LH, DU, PC

Data analysis and visualization: AP, LH, AG

Figures: AP, PG, LH

Testing and validation: AP, LH, AG

Supervision: SG, DU, RK, LG, OP, TM, MT

Writing—original draft: AP, PC, DU, PG

Writing—review & editing: AP, SG

