## Supplementary Figures for "Transformer-based spatial-temporal detection of apoptotic cell death in live-cell imaging"

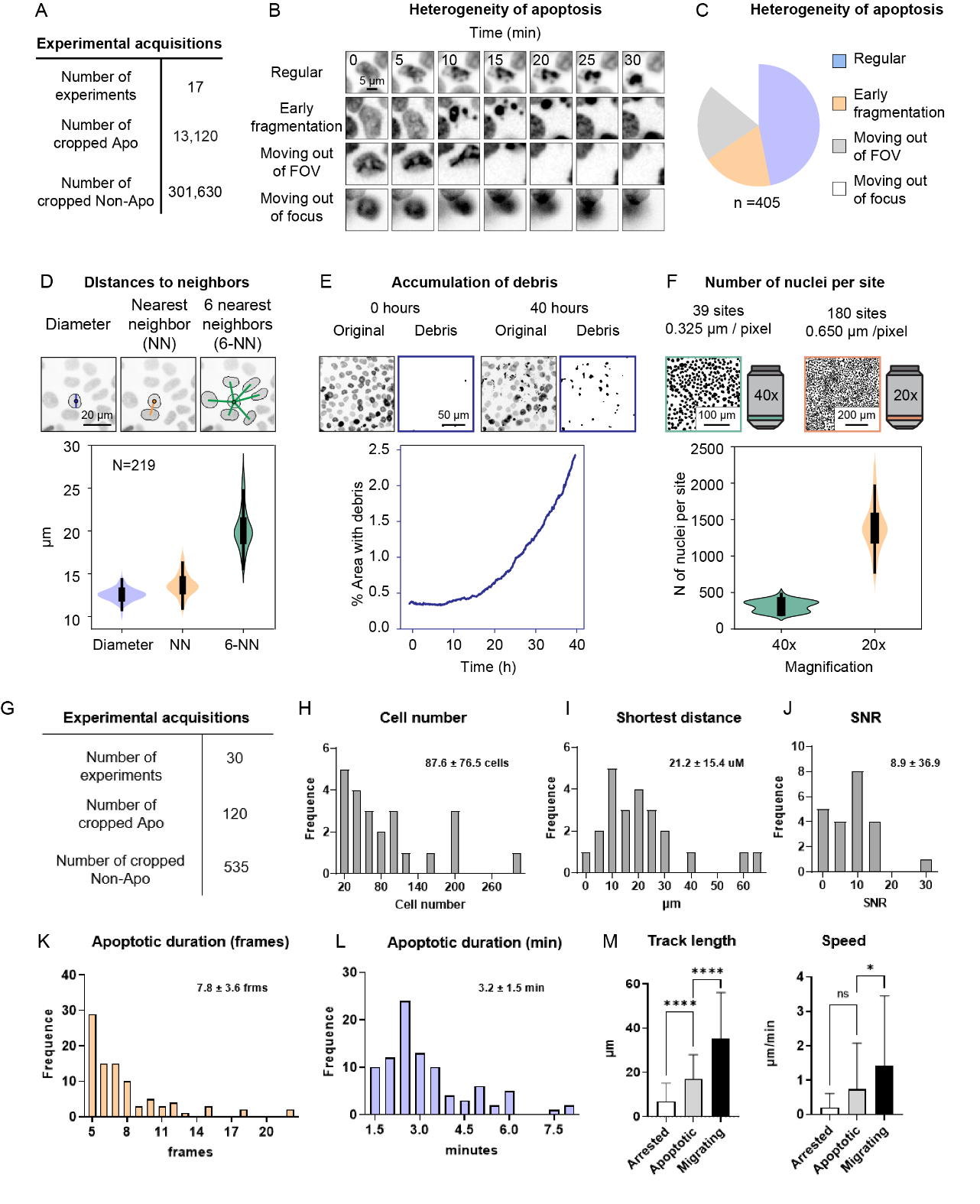

***Supplementary figure 1. Generation of* in vitro *and* in vivo *microscopy data. A.*** *Table reporting the number of entries of the dataset* in vitro*.* ***B.*** *Time-lapses showing the heterogeneity of the morphological appearance of apoptotic events.* ***C.*** *Pie chart representing the frequency of the classes of morphological appearance in the entire dataset.* ***D.*** *The density of the epithelium is quantified by comparing the diameter of the nuclei versus the distance to the nearest neighbor (NN), and to the 6 nearest neighbors (6-NN). Violin plot showing the mean of all cells of the first frame from (n = 219 FOVs).* ***E.*** *Data from a single field-of-view (FOV) shows the accumulation of apoptotic debris over time, making the identification of newer apoptotic events difficult. In this experiment, MCF10A cells were treated with 1.25 µM Doxorubicin for 40 hours. The image crops show the original nuclear channel and the binary images with identification of debris with a machine learning approach (Ilastik) and thresholding. The Chart represents the area occupied by debris over time.* ***F.*** *Two imaging modalities were used (40x, 20x), representative nuclear masks are shown in the left images. Violin plots show the mean number of nuclei in the first frame per FOV (40x: n = 39, 20x: n = 180).* ***G.*** *Table reporting the number of entries of the dataset* in vivo*.* ***H-J.*** *Quantification of cell numbers, shortest distance, and signal to noise ratio (SNR) in the generated IV-2PM movies (n = 30).* ***K-L.*** *Histograms representing the duration of the apoptotic events expressed in frames (****K****) and minutes (****L****).* ***M.*** *Quantification of the track length (****left****) and cell speed (****right****) of apoptotic cells before disruption, compared to* *arrested and migrating cells. Statistical comparison was performed with non-parametric Kruskal-Wallis test. Columns and error bars represent the mean and standard deviation respectively. Significance is expressed as: p ≤* *0.05 (*), p ≤* *0.01 (**), p ≤* *0.001 (***), p ≤* *0.0001 (****).*

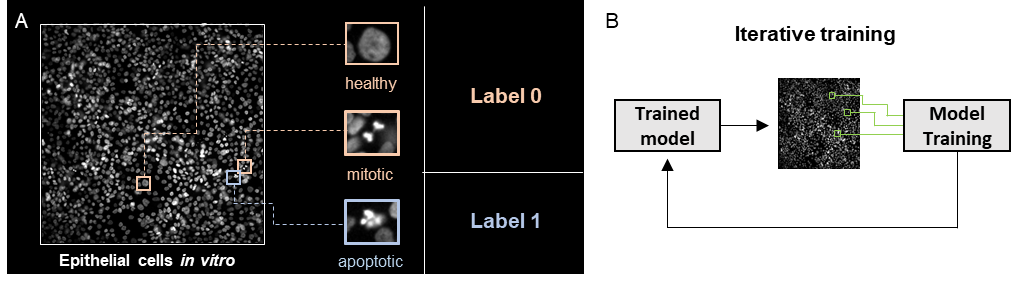

***Supplementary figure 2. Training and performance* in vitro*.*** ***A.*** *The training set describes a binary classification task in which the class label 1 contains the nuclei of epithelial cells undergoing apoptosis, while the class label 0 includes healthy and mitotic nuclei.* ***B.*** *The class label 0 has been further expanded by iteratively including false positives generated by a trained network and applied to movies which contained no apoptotic events*

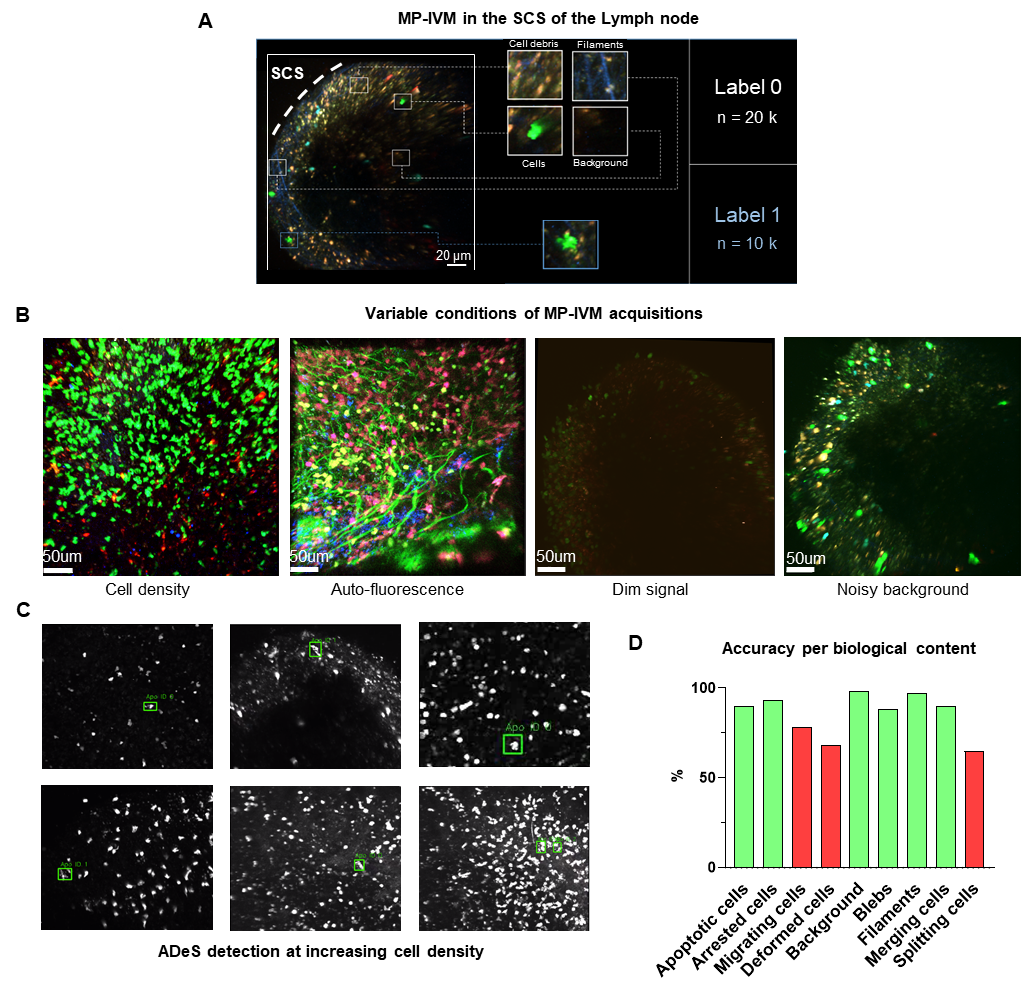
***Supplementary figure 3. Training and deployment* in vivo*. A.*** *The designed training set describes a binary classification task in which the class label 1 contains only apoptotic cells and the class label 0 encompasses all non-apoptotic content, including healthy cells, filaments, background, and cell debris.* ***B.*** *Representative snapshots of variable and potentially challenging conditions in multi-photon intravital microscopy (MP-IVM), including high cell density, auto-fluorescence, dim signal, and noisy background.* ***C.*** *Representative micrographs depicting the detection of apoptotic cells at increasing cell densities.* ***D.*** *Graph showing the accuracy of ADeS in predicting the class label (0 or 1) of sequences containing different biological content. Red bars represent an accuracy below 80%.*

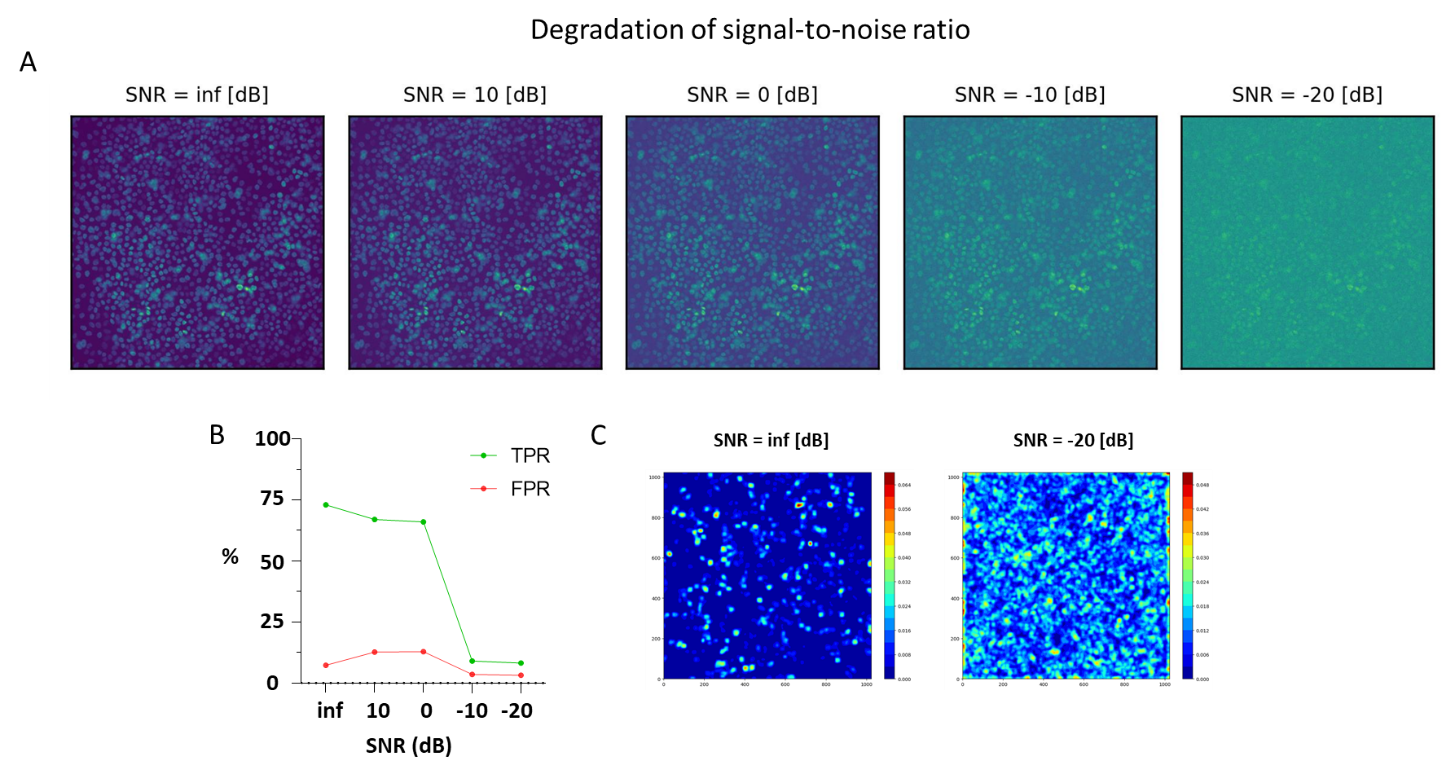

***Supplementary figure 4. Effect of noise on ADeS performance. A****.* In vitro *micrographs showing the same acquisition with the addition of different levels of noise measured in decibels (dB). High SNR values represent better video quality.* ***B****. Plot showing the effect of increasing noise levels on the true positive rate (TPR) and false positive rate (TPR) of ADeS predictions* ***C.*** *Noise effect visualized as a likelihood heatmaps. Higher noise generate frequent low confidence predictions.*

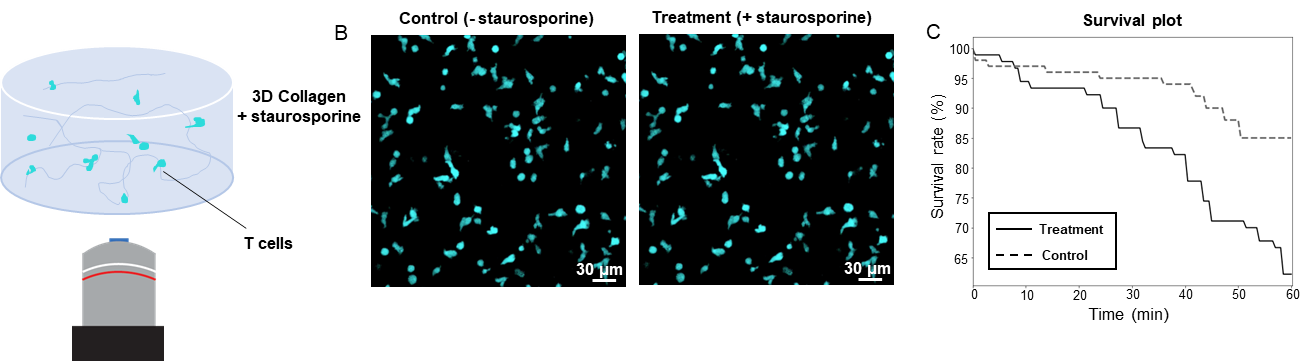

***Supplementary figure 5. Applications for toxicity assay* in vitro*. A****. Schematic drawing representing in vitro****-****cultured* *T cells treated with with staurosporine.* ***B****. Confocal micrograph snapshot showing T cells at 60 min after treatment with staurosporine (****left****) compared to the untreated control group (****right****).* ***C****. Survival assay plot of control (dotted lined) and treated samples (solid line) during the first 60 min post treatment with staurosporine.*
